# Aging-associated weakening of the action potential in fast-spiking interneurons in the human neocortex

**DOI:** 10.1101/2024.03.24.586453

**Authors:** Viktor Szegedi, Ádám Tiszlavicz, Szabina Furdan, Abdennour Douida, Emoke Bakos, Pal Barzo, Gabor Tamas, Attila Szucs, Karri Lamsa

## Abstract

Aging is associated with the slowdown of neuronal processing and cognitive performance in the brain; however, the exact cellular mechanisms behind this deterioration in humans are poorly elucidated. Recordings in human acute brain slices prepared from tissue resected during brain surgery enable the investigation of neuronal changes with age. Although neocortical fast-spiking cells are widely implicated in neuronal network activities underlying cognitive processes, they are vulnerable to neurodegeneration. Herein, we analyzed the electrical properties of 147 fast-spiking interneurons in neocortex samples resected in brain surgery from 106 patients aged 11–84 years. By studying the electrophysiological features of action potentials and passive membrane properties, we report that action potential overshoot significantly decreases and spike half-width increases with age. Moreover, the action potential maximum-rise speed (but not the repolarization speed or the afterhyperpolarization amplitude) significantly changed with age, suggesting a particular weakening of the sodium channel current generated in the soma. Cell passive membrane properties measured as the input resistance, membrane time constant, and cell capacitance remained unaffected by senescence. Thus, we conclude that the action potential in fast-spiking interneurons shows a significant weakening in the human neocortex with age. This may contribute to the deterioration of cortical functions by aging.

## 1. INTRODUCTION

Aging affects all body cells. The functions of body parts (including the brain) peak in young adulthood, after which they decline until they cease completely via death. Humans generally outlive other mammals, particularly rodents (which are the most commonly used laboratory animals in biomedical brain research). Neurons are highly differentiated postmitotic cells, and for most of them, the lifespan is equal to that of the entire organism (Aranda-Anzaldo and Dent, 2017; Herrup, 2013; Isaev et al., 2019). Therefore, to understand age-related changes in human cerebral neurons, it is essential for aging to be investigated using human brain tissue. There is limited access to human brain tissue to investigate neuronal functions; however, in recent years, it has become increasingly common to use human brain tissue resected during brain surgery, which often contains a small block of the neocortex in which a surgeon needs to make a hole to gain access to the pathological site in the deep brain area (Straehle et al., 2023). The neocortex is part of the telencephalon and is the site in the brain where the most complex brain functions occur (Higley and Cardin, 2022; Liu and Konopka, 2020; Luhmann, 2023). Resected neocortical tissue samples provide highly relevant material for human neuron aging investigation because noninvasive brain scan studies, behavioral tests, and anatomical postmortem analyses show that aging affects the neocortex in various ways (Peters, 2006; Rozycka and Liguz-Lecznar, 2017; Sibille, 2013). However, to the best of our knowledge, only a few studies have investigated electrophysiological changes in neurons in the human brain during senescence (Guet-McCreight et al., 2023; Pegasiou et al., 2020; Popov et al., 2023; Testa-Silva et al., 2014; Wang et al., 2016).

Animal experiments have shown that aging-induced changes in neurons develop in different ways and at different rates in distinct types of neurons in the brain (Radulescu et al., 2021; Rozycka and Liguz-Lecznar, 2017), similar to various neurodegenerative diseases where some nerve cell types manifest functional changes more sensitively than others (Temple, 2023). The high metabolic demand of fast-spiking interneurons in the neocortex predisposes them to aging-associated brain changes, and it is expected that these neurons will be among the first to suffer from blood flow compromised in the aged brain (Beishon et al., 2021; Cunnane et al., 2020; Ibrahim and Llano, 2019; Kann, 2016). Fast-spiking interneurons play a central role as organizers of the time and space of principal neurons’ activity in the cortex; consequently, these inhibitory cells orchestrate neuronal network activities underlying complex sensory processes, cognitive tasks, and learning (Druga et al., 2023; Hu et al., 2014). Rodent experiments have shown that cognitive and sensory decline in many neurodegenerative diseases is associated with the altered function of fast-spiking interneurons in the neocortex (Cernotova et al., 2023; Hijazi et al., 2023), and various studies have also reported age-associated changes in these neurons (Dong et al., 2023; Lehmann et al., 2012; Mattson, 2020). However, little is known about the functional changes in these delicate nerve cells during aging in the neocortex of the human brain.

Herein, we study fast-spiking neurons in human acute brain slices prepared from neocortical tissues resected during deep brain surgery. We focus on two key electrophysiological features of neurons, which are the strength and kinetics of action potentials and the passive membrane properties underlying neuron’s intrinsic electrical excitability.

Our data demonstrate aging-induced action potential attenuation, whereas parameters monitoring neurons’ passive electrical properties remain unchanged from youth to old age. We observed age-associated alterations in the action potential rise speed, duration of the depolarization phase, and peak depolarizing membrane potential value. These changes indicate a reduction in the action potential sodium current generated in the neuron’s soma or proximal dendrite. We suggest that these changes may partially underlie aging-associated slowdown of computational processes and impaired learning in the neocortex of the human brain.

## 2. MATERIALS AND METHODS

### 2.1 Ethics statement

All procedures were approved by the University of Szeged Ethics Committee and Regional Human Investigation Review Board (ref. 75/2014), and they were conducted per the tenets of the Declaration of Helsinki. Written informed consent for the study of excised human cortical tissue was obtained from all patients before surgery.

### 2.2 Human brain slices

Neocortical slices were prepared from samples of the frontal, cortex, temporal cortex, or other cortical areas (Fig. 1A) removed for the surgical treatment of deep brain targets. Samples were acquired from the left and right hemispheres of males and females (Fig. 1B) operated as treatment for tumor or neoplasia, malformation (such as cyst or aneurysm), or increased intracortical pressure (Fig. 1C). Anesthesia was induced with intravenous midazolam (0.03 mg/kg) and fentanyl (1–2 mg/kg) following a bolus intravenous injection of propofol (1–2 mg/kg). The patients also received 0.5 mg/kg rocuronium to facilitate endotracheal intubation. Patients were ventilated with a 1:2 O_2_/N_2_O mixture during surgery, and anesthesia was maintained with sevoflurane. The resected tissue blocks were immersed in an ice-cold solution containing 130 mM NaCl, 3.5 mM KCl, 1 mM NaH_2_PO_4_, 24 mM NaHCO_3_, 1 mM CaCl_2_, 3 mM MgSO_4_, and 10 mM D(+)-glucose aerated with 95% O_2_/5% CO_2_ within a glass container. The container was then placed on ice inside a thermally isolated box and transported from the operating room to the electrophysiology laboratory with continuous 95% O_2_/5% CO_2_ aeration. Slices measuring 350 μm in thickness were prepared from the tissue block using a vibrating blade microtome (Microm HM 650 V) and then incubated at 22°C–24°C for 1 h in slicing solutions. The slicing solution was gradually replaced with a recording solution (180 mL) using a pump (6–8 mL/min). The contents of the recording solution were identical to those of the slicing solution, except that it contained 3 mM CaCl_2_ and 1.5 mM MgSO_4_.

**Figure 1.**
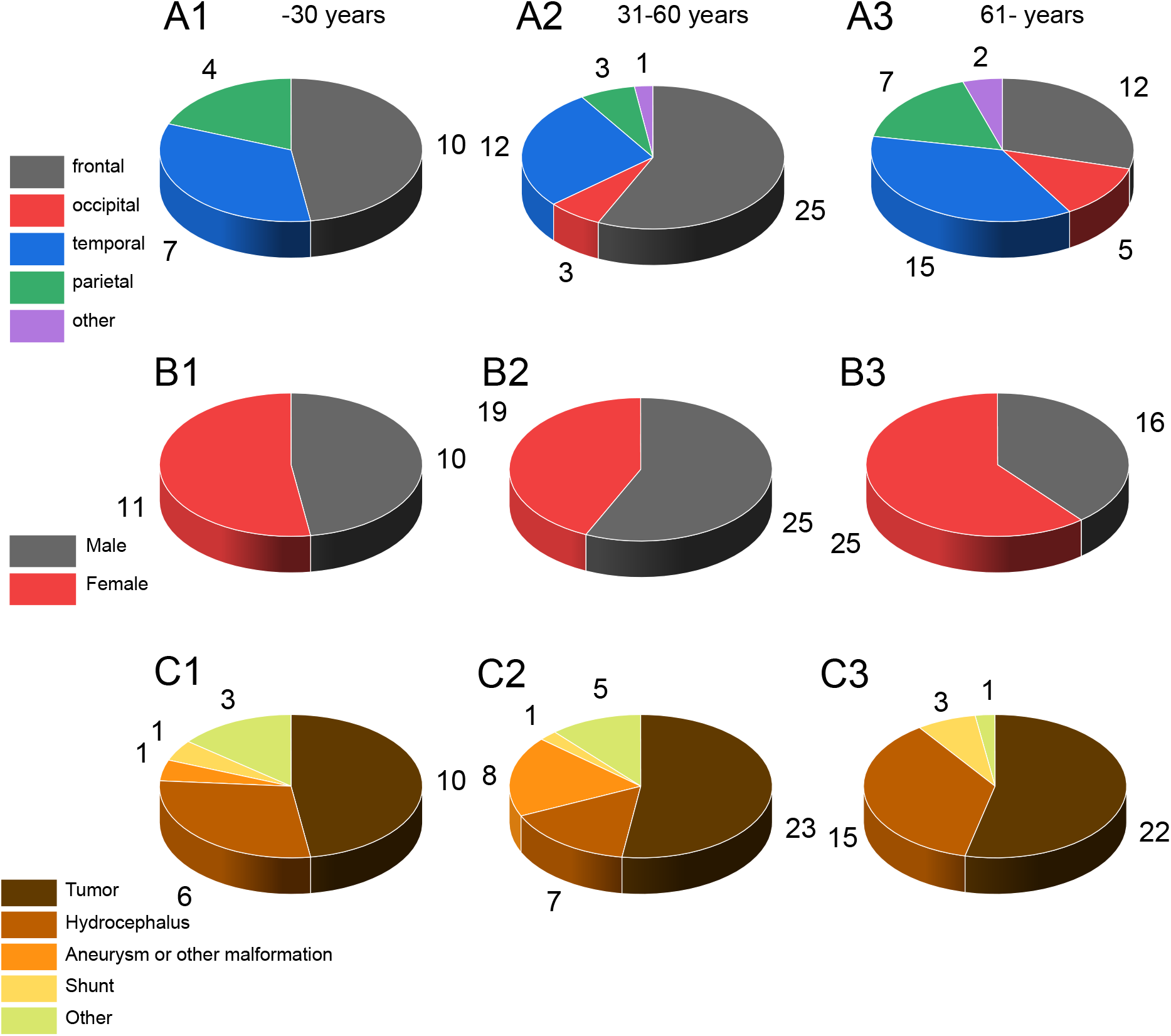
Proportions of resected brain tissue from different neocortical areas, patient sex, and primary diagnosis. A. The proportion of resected brain tissues from different neocortical areas used in the study In the three different age categories. A1) Fast-spiking cells recorded in tissue samples taken from patients aged ≤30 years. A2) Cells recorded in tissue from patients aged 31–60 years. A3) Cells in tissue from patients aged ≥61 years. Distinct neocortical areas (frontal, occipital, temporal, parietal, or other cells with no information) are color-coded in pie charts, as indicated in the inset. B. Patient sex in the three age groups B1–B3. C. Primary categorical diagnosis for surgery in the three age groups C1–C3.

### 2.3 Electrophysiology

Recordings were performed in a submerged chamber perfused at 8 mL/min with the recording solution maintained at 36°C–37°C. Cells were patched under visual guidance using infrared differential interference contrast video microscopy and a water-immersion 20× objective with additional zoom (up to 4×). Micropipettes (5–8 MΩ) for whole-cell patch-clamp recording were filled with intracellular solution containing: 126 mM K-gluconate, 8 mM NaCl, 4 mM ATP-Mg, 0.3 mM Na_2_-GTP, 10 mM HEPES, and 10 mM phosphocreatine (pH 7.0–7.2; 300 mOsm) and supplemented with 0.3% (w/v) biocytin for subsequent staining with Alexa488- or Cy3-conjugated streptavidin. Recordings were performed using a Multiclamp 700B amplifier (Axon Instruments) and low-pass filtered at a 6–8 kHz cutoff frequency (Bessel filter). The series resistance and pipette capacitance were compensated online. Membrane potential values were not corrected for the liquid junction potential error. Somatic input resistance, membrane time constant, and capacitance were measured in the current-clamp mode by applying current (250–500 ms) to hyperpolarize the membrane to −90 mV from −70 mV. All parameters were measured from at least five traces. The intrinsic property step protocol was systematically applied to 113 of the 147 fast-spiking neurons.

### 2.4 Fast-spiking neuron identification

Cells were primarily identified by narrow action potential (action potential half-width <0.5 ms, n = 79 cells). In addition, if the spike width exceeded 0.5 ms, the cell was considered a fast-spiking interneuron if it showed high-frequency firing (>150 Hz) with modest firing frequency accommodation (n = 68 cells) (Petilla Interneuron Nomenclature et al., 2008). Spiking frequency accommodation was measured during a 500 ms square-pulse depolarizing step that elicited consistent >150 Hz firing in the cells. Frequency adaptation was measured at 400–500 ms period of consistent firing and compared with firing frequency at the initial 100 ms. The median adaptation value was 0.900, with lower and upper quartiles of 0.850 and 0.933, respectively (Petilla Interneuron Nomenclature et al., 2008).

We tested 33 of the 79 cells in the first category, and 41 of the 68 cells in the second category for parvalbumin (pv) through immunohistochemistry; they were all immunopositive for pv. Immunoreactions and analyses were performed as described by Szegedi et al. (2023).

### 2.4 Single-cell model

The three-compartmental model neuron, including its cylindrical dendritic and axonal compartments, was based on the formalism described previously (Szegedi et al., 2023). The model consists of a somatic compartment, one axonal compartment, and one dendritic cable compartment, the axon being 140 µm long and measuring 0.8 µm in diameter and the dendrite measuring 200 µm in length and 1 µm in diameter. Cable parameters were as follows: axon longitudinal resistance = 0.7 MΩ*µm; axon capacitance = 6 fF/µm^2^. The dendrite (also referred to here as the “primary dendrite”) has twice the capacitance of the axon, 12 fF/µm^2^. The ohmic resistance of the somatic compartment was 700 MΩ, and the capacitance of the somatic compartment was 25 pF. The leakage reversal potential was set to −68 mV. The model incorporated six voltage-dependent currents: one high-threshold and one low-threshold transient Na current, a delayed rectifying K current (Kd), the HCN current, the slowly inactivating D-type K current, and the inward rectifying K current (Kir). The voltage-dependent Na, Kd, and D currents were assigned to the axonal and somatic compartments, whereas the HCN and Kir currents were included in the somatic and dendritic compartments. Parameters such as the rheobase and voltage sag were chosen to replicate basic response properties and carefully adjusted using experimental data to attain the best match between simulated and recorded voltage traces. All intrinsic voltage-dependent currents were calculated using the following equation:

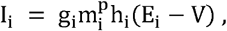

where i represents the individual current type, g_i_ is the maximal conductance of the current, m_i_ is the activation variable, p is the exponent of the activation term, h_i_ is the inactivation variable (either first-order or absent), and E_i_ is the reversal potential. The activation (m) and inactivation (h) kinetics were modeled according to the following equation:

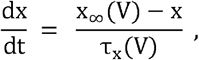

where x represents m or h, and voltage-dependent steady-state activation and inactivation are described by the sigmoid function

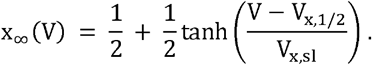

The midpoint V_x,1/2_ and slope V_x,sl_ parameters of the sigmoids, and the other kinetic parameters are shown in Table 1. The time constants (*τ*) of activation and inactivation were modeled as bell-shaped functions of the membrane potential (V) according to the following equation:

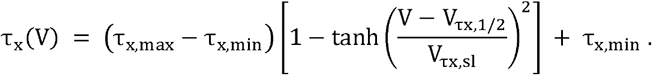

### 2.5 Data analysis

Data were acquired using Clampex software (Axon Instruments) and digitized at 35–50 kHz. The data were analyzed offline using pClamp (Axon Instruments), Spike2 (version 8.1, Cambridge Electronic Design), OriginPro (OriginLab Corporation), and SigmaPlot14 software (Systat Software).

#### Statistics

Results are presented as medians with interquartile ranges. Comparisons between variables were performed using the ANOVA on ranks with Dunn’s post hoc pairwise test, and Spearman Rank Order correlation. Splitting into age categories was necessary to get similarly-sized age groups while using standard age steps (30 years).

## 3. RESULTS

To study changes occurring during aging in human neocortical neurons, we analyzed 147 fast-spiking interneurons recorded with whole-cell patch clamps in acute brain slices of neocortical tissue resected during brain surgery from 106 patients. We report fast-spiking cell parameters as the mean values per patient, as well as the values per individual cell. All tissue came from deep brain surgery, where neocortical tissue was resected to allow access to the deep brain pathological locus. Fast-spiking neurons recorded in layer 2/3 of the frontal, temporal, parietal, and occipital areas (Fig. 1) were identified by their short action potential duration (mean half-width: 0.557 ± 0.156 ms) with weak spiking frequency accommodation. Cells were identified as fast-spiking interneurons by showing a narrow action potential (action potential half-width < 0.5 ms, n = 79 cells) or if the spike width was wider than 0.5 ms, by modest firing frequency accommodation and high-frequency firing capacity (n = 68 cell) (see Methods). We tested 74 of the 147 cells for parvalbumin, a marker protein for fast-spiking interneurons, through immunohistochemistry; all cells were immunopositive. This study focuses on two key electrophysiological parameters: action potential and passive electrical membrane properties underlying the intrinsic excitability of neurons. The relative proportion of tissue from neocortical areas of the brain, the patient sex and the primary diagnosis are shown in Fig. 1A-C.

### 3.1 Action potentials become weaker during aging

We observed a significant reduction by age of the action potential (AP) overshoot; i.e., the peak of the AP positive deflection. The AP was evoked by a short (250 ms) depolarizing step above the firing threshold at a membrane potential of −70 mV. We determined the lowest depolarizing step triggering AP using incrementally increasing positive current steps (step-wise increments from 10 pA to 20 pA) and systematically analyzed the first AP evoked by depolarization.

When analyzing the action potential’s positive peak value (see Fig. 2A1) measured as the average of cells per patient we found significant reductions by human age (n = 106 patients, r = −0.201, p = 0.0458, Spearman correlation) (Fig. 2A2). In addition, the AP positive peak value of individual cells was significantly correlated with age (n = 147, r = −0.1947, p = 0.0181, Spearman correlation) (Fig. 2A3). We compared AP overshoot data among age groups by dividing the average per patient data into three age groups (“≤30 years old,” “31– 60 years old,” and “≥61 years old”). This can reveal changes in parameters occurring only at young or old age. We found a significant difference between the youngest (≤30 years) and the oldest (≥61 years) group (n =21 and 41, respectively; p = 0.028, ANOVA on Ranks with Dunn’s post hoc pairwise test) (Fig. 2A4). The overshoot in the age groups was (as median and quartile range): 19.36 mV, 11.62–31.40 mV at “≤30 years” (n = 21 patients); 16.52 mV, 9.17–24.65 mV at “31–60 years” (n = 44 patients); 12.62 mV, 5.88–18.39 mV at “≥61 years” (n = 41 patients).

**Figure 2.**
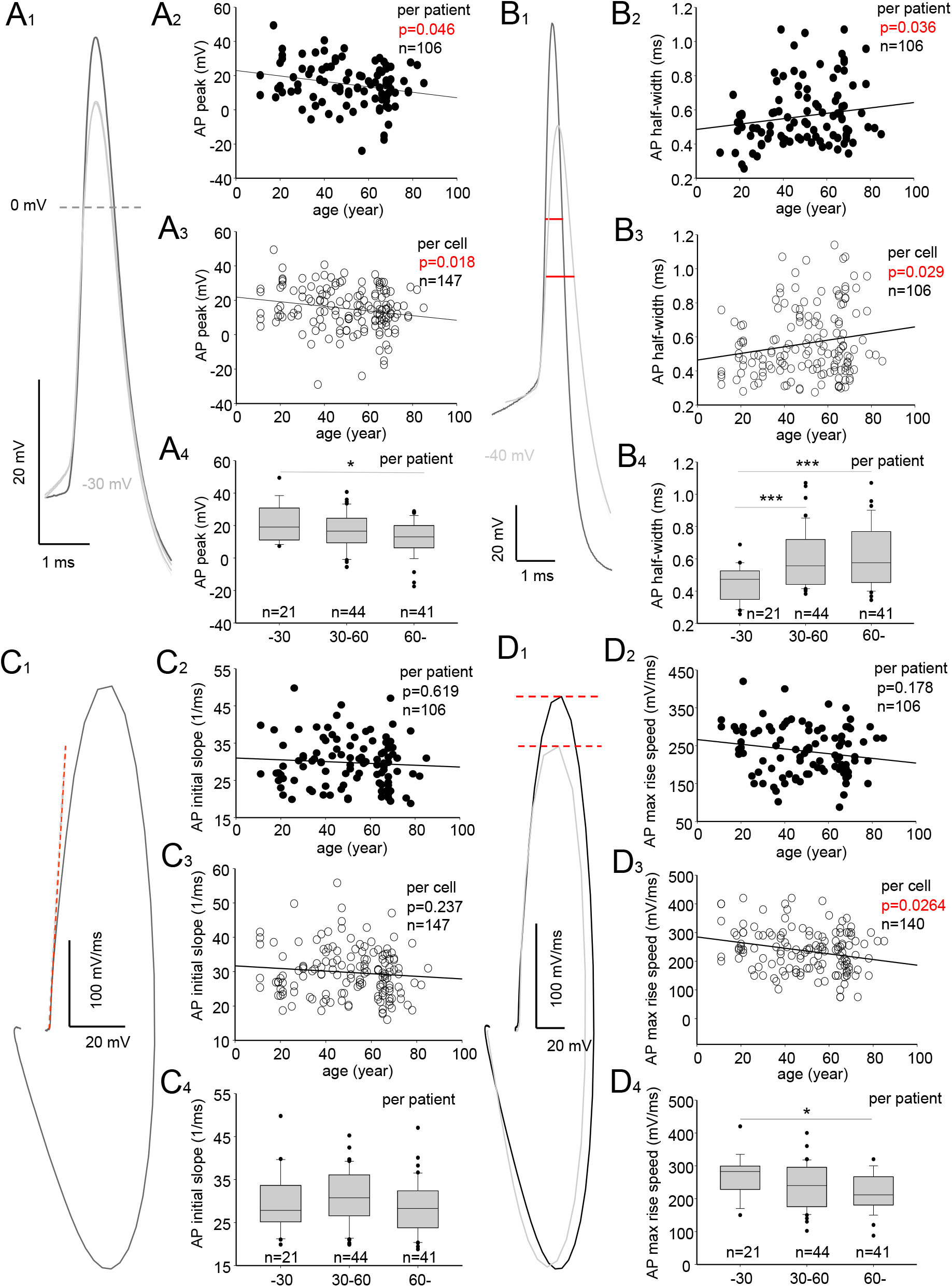
The action potential becomes weaker and prolonged with age in human fast-spiking interneurons. A. The reduced positive peak membrane potential value by action potential in patients of different ages. A1) Sample action potential traces in fast-spiking cells representing median positive peak potential values observed in the different age categories: ≤30 years (black) and 61 years or older (gray). Traces are averages of 5. A2) Graph showing a negative correlation between peak positive membrane potential value the action potential reaches (AP peak) and human age in 106 patients (Spearman correlation). Each AP peak value represents the average in a single patient. A3). Negative correlation between AP peak and human age for individual cell values (n = 147) (Spearman rank correlation test). A4) Box plots show median, upper, and lower quartile values, minimum and maximum (whiskers), and outliers (dots) for AP peak (average value in patient). There was a significant difference between the youngest (≤30 years) and oldest (≥61 years) groups (ANOVA on Ranks with Dunn’s post hoc pairwise test). B. Widening of the action potential by age. B1) Sample traces showing the half-width of the positive membrane potential deflection of the action potential. Traces show action potentials observed in the age categories “≤30 years” (black) and “≥61 years” (gray). Traces are averages of five. B2) Positive correlation between the half-width and human age in 106 patients (Spearman correlation). Each value represents the average in a single patient. B3) Positive correlation between the half-width in individual cells (n = 147) and human age (Spearman rank correlation test). B4) The box plots show the median and interquartile range, minimum and maximum (whiskers), and outliers (dots) for half-width average in patients. The two oldest age groups showed significantly longer half-width values than the youngest group (ANOVA on ranks, Dunn’s post hoc pairwise test). C. The action potential initial phase is unchanged. C1) Sample trace of the action potential phase plot showing its first derivative (dV/dt) against the membrane potential (Vm). Dotted red line illustrates the maximum slope of the tangent line in the phase plot above the spike threshold level of 10 mV/ms. This defines the phase slope (ms^−1^) of AP onset, which is used to measure the action potential initial phase (which reflects the sodium channel activity generated in the axon’s initial segment) (see Methods for details). C2) The graph shows the absence of a correlation between the action potential initial rise slope value average in patients and human age in 106 patients (Spearman correlation). C3) Absence of a correlation between the initial rise slope of 147 individual cells and human age (Spearman correlation). C4). Comparison of the initial rise slope values (average in patient) between the three age groups (ANOVA on ranks). D. Action potential maximum-rise speed decreases from young to old age. D1) Sample phase plots (dV/dt against Vm) of the action potential showing maximum-rise speed values (indicated with red dotted lines) in cells at young “≤30 years” (black) and old “≥61 years” age (gray). Traces are averages of five. The maximum-rise speed mainly reflects soma-generated sodium channel activity. D2) Although the action potential maximum-rise speed average in patients failed to show correlation with human age (Spearman correlation), there was a strong negative correlation between the action potential maximum-rise speed in individual cells and human age (Spearman correlation) (D3), and significant difference in the maximum-rise speed value between the youngest and oldest age groups. Plot shows differences between average value per patient in “≤30 years” and “≥61 years” age groups (ANOVA on Ranks with Dunn’s post hoc pairwise test).

Action potential also becomes slower during aging. We found that half-width, measured as the width of the AP depolarization phase at the mid-amplitude point between the resting potential and peak amplitude (Fig. 2B1), significantly increased with age. The average AP half-width per patient and age showed a positive correlation (n = 106, r = 0216, p = 0.0311, Spearman correlation) (Fig. 2B2). In addition, the AP half-width of individual cell per age showed correlation (n = 147, r = 0.1781, p = 0.0287, Spearman correlation) (Fig.2B3). The average AP half-width per patient in the youngest age group (up to 30 years) was significantly different from the mid-age group (p = 0.009), as well as from the oldest (61 years and older) group (p = 0.006) (ANOVA on Ranks with Dunn’s post hoc pairwise test) (Fig. 2B4). The half-width values in the three age groups were (median and quartile range): 0.473 ms, 0.348–0.526 ms at “≤30 years” (n = 21 patients); 0.556 ms, 0.441–0.720 ms at “31–60 years” (n = 44 patients); 0.575 ms, 0.452–0.769 ms at “≥61 years” (n = 41 patients).

Because the altered variables above indicate a weakening of the AP depolarization phase, we also analyzed the AP rise speed and rise slope. The initial slope and rise phase maximum speed are shown in Fig. 2C-D. The two parameters reflect sodium channel activity in the axon (axon initial segment) and in the soma with the proximal dendritic area, respectively. We found that the axon potential rise phase initial slope (Fig.2C1) was not significantly changed by aging. The rise phase initial slope was defined as the maximum slope of the tangent line in the AP phase plot above a spike threshold level of 10 mV/ms. The tangent line provides the AP onset initial rapidity (ms^−1^). The rise slope (average in patient) versus age did not show a correlation (n = 106, r = −0.0498, p = 0.619, Spearman correlation) (Fig. 2C2), nor did the rise slope of individual cell and human age did not show correlation either (n = 147, r = −0.0981, p = 0.2372, Spearman correlation) (Fig. 2C3). In addition, there was no difference in slope between the age groups (p = 0.261) (ANOVA on Ranks) (Fig.2C4). The rise phase initial slope values in the three age groups were (median and quartile range): 27.826 ms^−1^, 25.101–33.688 ms^−1^ at “≤30 years” (n = 21 patients); 30.771 ms^−1^, 26.519–36.143 ms^−1^ at “31–60 years” (n = 44 patients); and 28.264 ms^-1^, 23.710–32.439 ms^-1^ at “≥61 years” (n = 41 patients). Thus, the AP rise phase component emerging from the axon, and all sodium channel activity was not affected by age.

The maximum speed of the AP rise phase (Fig. 2D1) failed to show any difference with age for the maximum-rise speed per patient (n = 106, r = −0.136, p = 0.178) (Fig. 2D2). However, the maximum-rise speed for individual cells significantly weakened with aging (n = 140, r = −0.188, p = 0.0264, spearman’s correlation) (Fig. 2D3). In addition, there was a statistically significant difference in the maximum-rise speed between the youngest and oldest age groups (p = 0.014) (ANOVA on Ranks with Dunn’s post hoc pairwise test) (Fig. 2D4). The maximum-rise speed in the three age groups was (median and quartile range) 283 mV/ms, 229–300 mV/ms at “≤30 years” (n = 21 patients); 240 mV/ms, 175–296 mV/ms at “31–60 years” (n = 44 patients); and 210 mV/ms, 180–260 mV/ms at “≥61 years” (n = 41 patients). The results show that between the youngest and oldest age group, there is reduced strength of AP rise speed, indicating a weakened sodium channel-mediated component of AP in the soma.

### 3.2 The AP afterhyperpolarization remains unaltered by age

Next, we investigated the AP repolarizing phase and after hyperpolarization (AHP) to determine whether these were also affected by age. However, unlike the AP depolarization phase, the repolarizing phase was not significantly changed during aging. There was no statistically significant relationship between age and the maximum speed of membrane potential change during the AP repolarizing phase (Fig. 3A1) measured as the average value per patient (n = 106, r = 0.0719, p = 0.476) (Fig. 3A2), or as individual cell value (n = 147, r = 0.1186, p = 0.155) (Spearman correlation) (Fig. 3A3). In line with this, there was no significant difference in the maximum repolarization speed between the three age groups (p = 0.337) (ANOVA on Ranks) (Fig. 3A4). The maximum repolarization speed was (median and quartile range) −150 mV/ms, −123 to −174 mV/ms at “≤30 years” (n = 21 patients); −140 mV/ms, −90 to −171 mV/ms at “31–60 years” (n = 44 patients); −128 mV/ms, −93 to −159 mV/ms at “≥61 years” (n = 41 patients).

**Figure 3.**
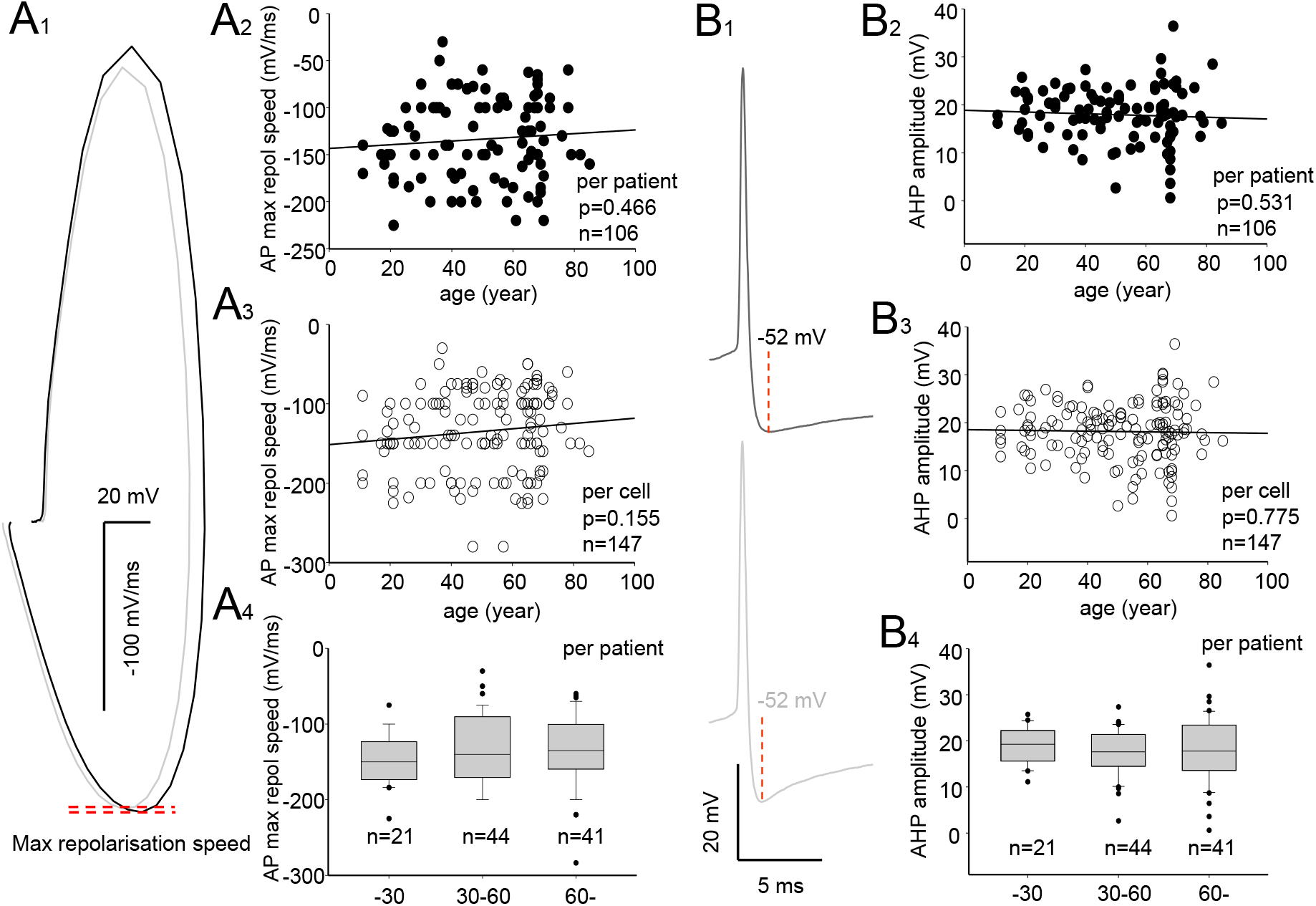
The action potential repolarization remains unchanged with age. A. The maximum repolarization speed of the action potential does not change with age. A1) Sample phase plots (dV/dt against Vm) of the action potential showing the maximum repolarization speed value (dotted red lines) in cells in the “≤30 years” (black) and “≥61 years” (gray) age groups. The maximum descending speed mainly reflects somatic and axonal potassium channel activity. A2) The maximum repolarization speed of the action potential (average in patient) in 106 patients did not show significant correlations with human age (Spearman correlation). A3) The maximum repolarization speed in 147 individual cells did not show a significant correlation with human age (Spearman correlation). A4) The three age groups did not differ significantly regarding the maximum repolarization speed (average value in patient in 106 patients) (ANOVA on ranks). B. The fast-spiking cell action potential afterhyperpolarization peak amplitude is not altered by human age. B1) Traces show the action potential AHP (amplitude indicated with a dotted line) in fast-spiking cells in the “≤30 years” (black) and “≥61 years” age categories. Traces are averages of five. B2) There was no significant relationship between AHP amplitude (average in patient) and human age in 106 patients (Spearman correlation). B3) There was no correlation between the AHP amplitude in individual cells (n = 147) and human age (Spearman correlation). B4) Comparisons of the AHP amplitude (average in patient) among the three age groups failed to show a statistically significant difference (ANOVA on Ranks).

Afterhyperpolarization, the amplitude measured from the onset of the AP rise phase to afterhyperpolarization peak negative value (Fig. 3B1) was also unchanged by aging. Correlation analyses failed to show a relationship between age and AHP amplitude measured as average per patient (n = 106, r = −0.0629, p = 0531) (Fig. 3B2), or as individual cell values (n = 147, r = 0.0238, p = 0.7752 (Fig. 3B3) (Spearman correlation). Comparisons of the AHP amplitude among the three age groups did not show significant differences (p = 0.766) (ANOVA on Ranks) (Fig. 3B4). AHP amplitude values (average per patient) in the groups were (median and quartile range) 19.27 mV, 15.53–22.31 mV at “≤30 years” (n = 21 patients); 17.61 mV, 14.42–21.46 mV at “31–60 years” (n = 44 patients); and 17.79 mV, 13.51–23.49 mV at “≥61 years” (n = 41 patients).

### 3.3 Passive electrical properties underlying the intrinsic excitability of neurons remain unaltered during aging

Given the significant changes in AP observed in fast-spiking interneurons, we also investigated whether the major passive electrical properties underlying the neuron’s electrical reactivity were affected by age. However, neurons showed no age-associated difference in cell soma capacitance, soma input resistance, or soma membrane time constant measured in whole-cell current clamps in fast-spiking cells.

There was no significant correlation between human age and cell capacitance measured as the average per patient (n = 81, r = 0.0828, p = 0.470) (Fig. 4A1), or as individual cell value (n = 133, r = 0.0096 and p = 0.919) (Fig. 4A2) (Spearman correlation). Also, there was no significant difference in capacitance among the three age groups (p = 0.738) (ANOVA on Ranks) (Fig. 4A3). Capacitance values (average per patient) in the age groups were (median and quartile range) 52.40 pF, 42.36–60.60 pF at “≤30 years” (n = 18 patients); 48.00 pF, 40.52–64.82 pF at “31–60 years” (n = 32 patients); and 51.69 pF, 43.65–72.61 pF at “≥61 years” (n = 31 patients).

**Figure 4.**
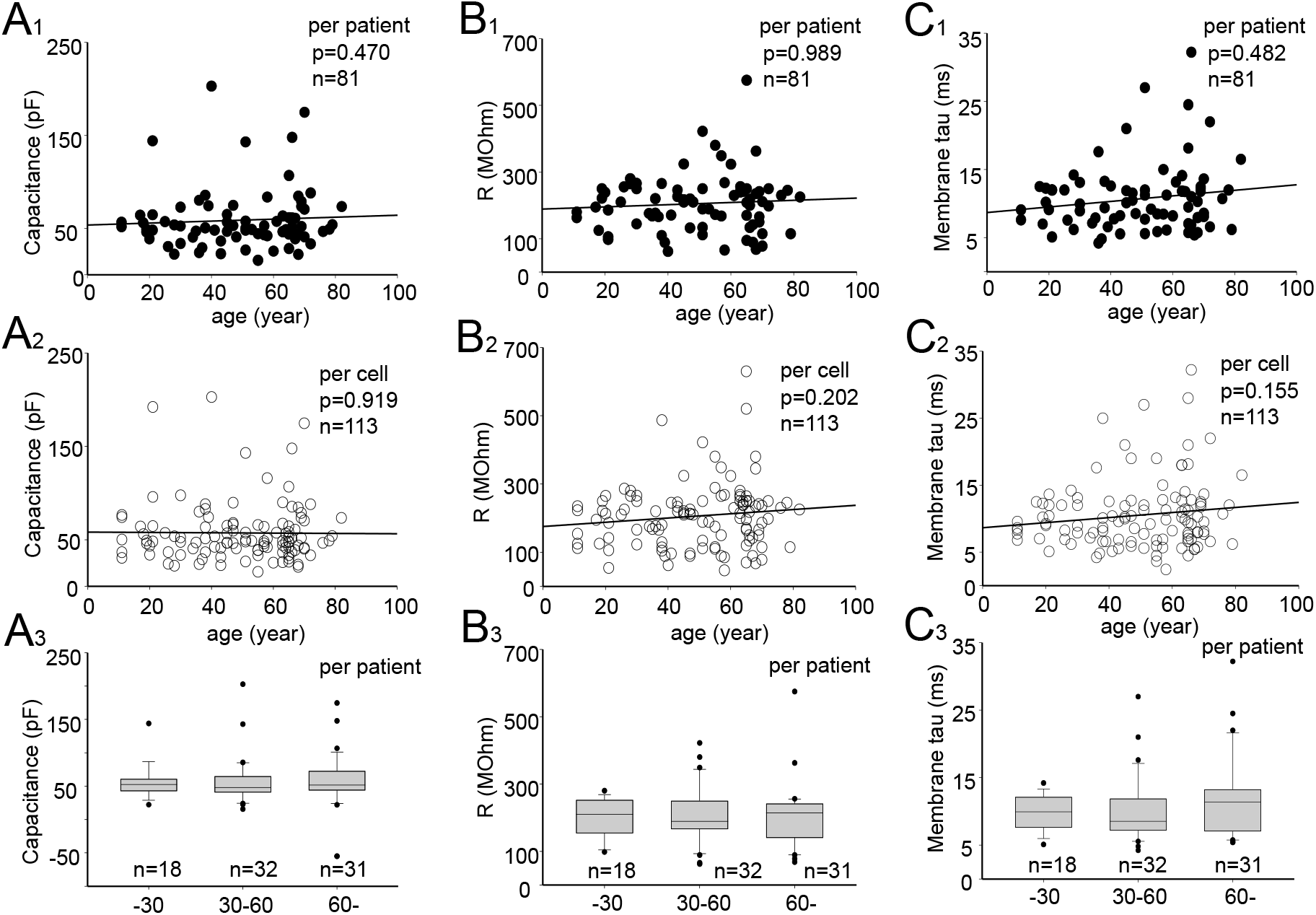
The passive electrical properties of fast-spiking cells remain unchanged during senescence. A. Similar cell capacitance values with age indicate similar soma sizes of recorded neurons. The graph shows the absence of a statistically significant correlation between human age and cell capacitance in fast-spiking cells measured as the average in patient (A1), or per individual cell value (A2) (Spearman correlation). A3) Comparisons of capacitance (average value in patient) between the three age groups did not show a significant difference (ANOVA on ranks). B. Similar input resistance in fast-spiking cells of different ages reflects unchanged passive membrane potential responsiveness to currents. B1) There was no significant correlation between cell input resistance (average value in patient) in 81 patients and human age (Spearman correlation). B2) Lack of significant correlation between cell input resistance in 113 individual cells and human age (Spearman correlation). B3) Comparisons of cell input resistance (average in patient) among the three age groups did not show a significant difference (ANOVA on ranks). C. Similar membrane time constant values with age demonstrate unchanged temporal reactivity of the cell membrane potential with age. C1) The membrane time constant (average in patient) in fast-spiking cells does not show a significant correlation in 81 patients with human age (Spearman correlation). C2) Lack of correlation with human age in 113 cells (Spearman correlation). C3) Comparison of membrane time constants (average in patient) among the three age groups did not show a significant difference (ANOVA on ranks).

There was no significant correlation between human age and input resistance measured as average per patient (n = 81, r = 0.0016, p = 0.989) (Fig. 4B1), or as individual cell value (n = 133, r = 0.1208 and p = 0.2024) (Fig. 4B2) (Spearman correlation). There was no statistically significant difference in resistance among age groups (p = 0.921) (ANOVA on Ranks) (Fig. 4B3). Input resistance values in age groups were as follows (median and quartile range): 210 MOhm, 154–253 MOhm at “≤30 years” (n = 18 patients); 189 MOhm, 166–250 MOhm at “31–60 years” (n = 32 patients); and 215 MOhm, 140–242 MOhm at “≥61 years” (n = 31 patients). In line with these results, we found that the membrane time constant (measured in soma to hyperpolarizing voltage steps at −70 mV to −90 mV) was not significantly correlated with age when tested as average value per patient (n = 81, r = 0.081, p = 0.482) (Fig. 4C1), or as individual cell (n = 113, r = 0.1345, p = 0.155) (Fig. 4C2) (Spearman correlation). Membrane time constant values (average per patient) in the three age groups were as follows (median and quartile range): 9.95 ms, 7.60–12.13 ms at “≤30 years” (n = 18 patients); 8.54 ms, 7.13–11.90 ms at “31–60 years” (n = 32 patients); and 11.38 ms, 7.01– 13.27 ms at “≥61 years” (n = 31 patients). The values did not differ significantly among groups (p = 0.471) (ANOVA on Ranks) (Fig. 4C3).

### 3.4 A single-cell computational model demonstrates age-associated changes in AP properties through decreased sodium channel-mediated activity in the soma

We used a computational single-cell model based on realistic fast-spiking neuron electrophysiological parameters measured in the whole-cell mode to mimic the real shape of the cell’s AP and firing behavior (Fig. 5). We investigated how well a selective reduction of the soma-located high-threshold voltage-gated sodium channel current explains the AP shape and overshoot alterations observed in real neurons by aging. We generated AP firing by depolarizing the membrane potential, akin to real whole-cell recorded neurons, in a three-compartment neuron model comprising the soma, main dendrite, and axon, with the AP initiation site being the axon’s initial segment and Hodgin–Huxley type AP with voltage-sensitive axonal low-threshold Na^+^ current, high-threshold Na^+^ current at the soma and proximal dendrite (Na-soma), and AP K^+^ current, voltage-insensitive K^+^ leak current, the slowly inactivating D-type K^+^ current, and Kir-type K^+^ current (Table 1). Following the simulation of the real shape of the cell’s AP (Fig. 5A), we selectively reduced the high-threshold sodium channel current located at the soma and primary dendrite while keeping other parameters unaltered. We simulated the AP with five different conductance (g) values for Na-soma (gNa-soma) starting with gNa-soma = 6 µS mimicking the AP recorded in a real fast-spiking neuron in the neocortex of a 21-year-old human (male, parietal cortex, resected in deep brain tumor operation). We selectively reduced gNa-soma to 5 µS, 4, 3, and to 2µS (Fig. 5A). The effect of the reduced gNa-soma on the AP waveform is illustrated in Figure 5B, which shows the spike phase plot depicting the AP’s first derivative (dVm/dt) against the membrane potential (Vm). This caused a reduction in the AP’s maximum-rise speed from 305 mV/ms to 260, and 225, 175, and 122 mV/ms, respectively (Fig. 5C). In parallel, the AP half-width, which was measured from the AP’s positive deflection, increased from 0.56 ms to 0.65 ms (Fig. 5D).

**Figure 5.**
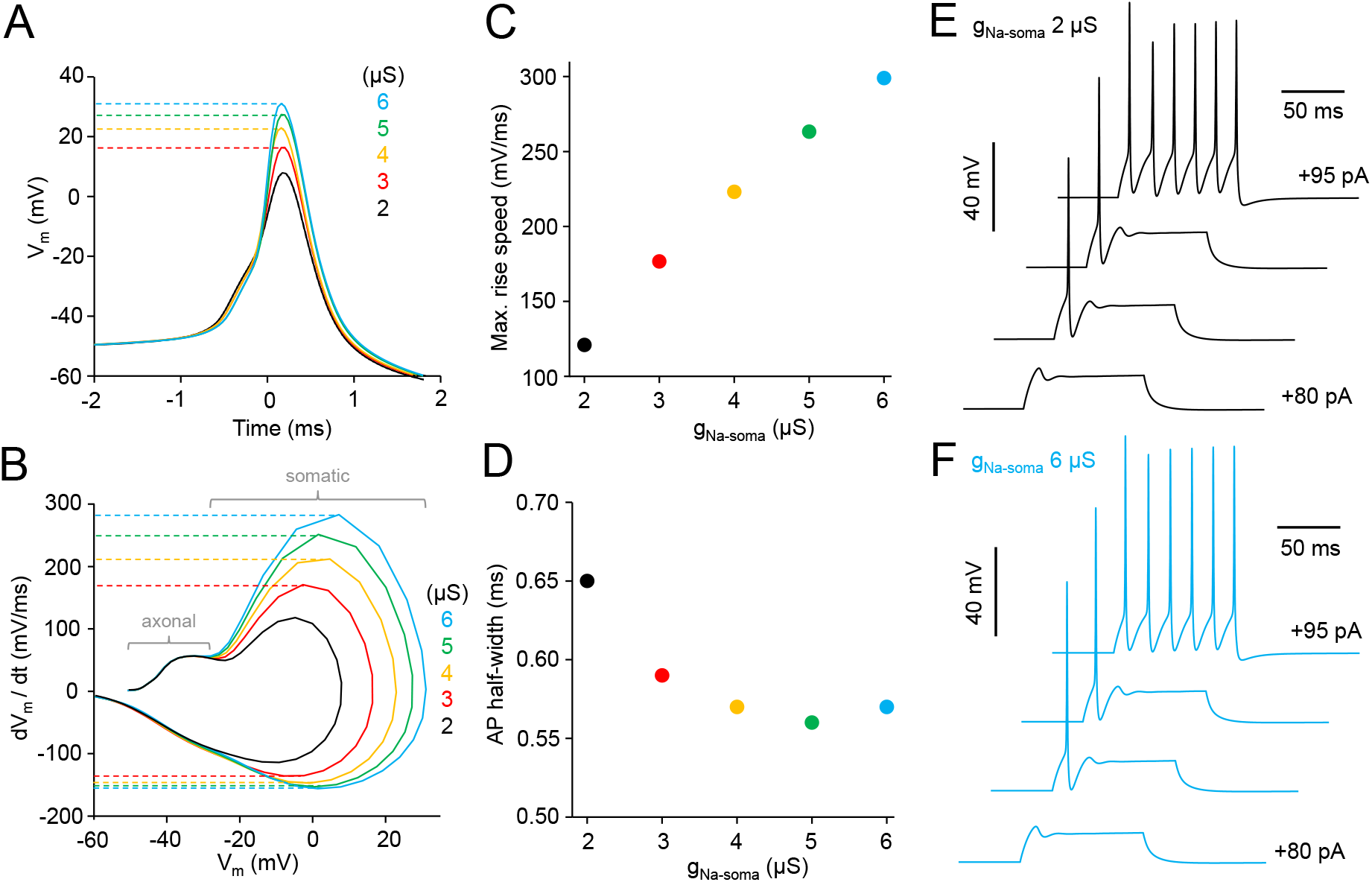
The simulation of a human fast-spiking neuron using a realistic single-cell computational model demonstrates AP widening and amplitude attenuation by reduced high-threshold sodium current in the soma. A single-cell model based on electrophysiological parameters recorded in a human fast-spiking neuron. A. Action potentials generated in the model cell. Superimposed spikes generated in model cells mimicking normal conditions with somatic sodium current conductance (gNa-soma) 6 µS on top (blue) and selectively reduced gNa-soma from 5 to 2 µS (color-coded from green to black). Dotted horizontal lines show the AP overshoot (positive peak amplitude) generated with different gNa values from 6 to 3 µS, and observed in real fast-spiking cells. B. Phase plot of the 5 simulated action potentials showing the action potential first derivative (membrane potential change speed as millivolts per ms, dVm/dt) as a function of the membrane potential (Vm). Dotted horizontal lines (color-coded by gNa strength) show that traces exhibit a pronounced reduction of its maximal rise speed (y-axis) from 305 mV/ms (with 6 µS) to 260 mV/ms (with 5 µS), 225 mV/ms (with 4 µS), 175 mV/ms (with 3 µS), and 122 mV/ms (with 2 µS). Concomitantly, the AP’s peak amplitude (X-axis) reduced with the three highest gNa-soma values from +31 mV to +27 mV and to +21 mV, respectively (not indicated with lines). The initial rise of the spike, generated in the axon remains unchanged. The maximum repolarization speed (dotted horizontal lines at the bottom) was only changed from −155 (with 6 µS) mV/ms to −151 mV/ms (5 µS) and to −147 mV/ms (4 µS) and to −143 mV/ms (3 µS). The axonal and somatic component of AP are indicated in the figure. C. The plot shows a positive correlation between the AP’s maximum-rise speed (mV/ms) in the simulated APs and the increasing gNa-soma conductance (X-axis) strength (P < 0.001) (Spearman correlation). D. This simulation shows that increasing the gNa-soma creates a shorter AP half-width time for positive deflection. E–F. The fast-spiking interneuron firing pattern generated by simulation with reduced 2 µS gNa-soma (E) and with 6 µS gNa-soma (F). Note the reduced AP overshoot in E compared to F.

The simulation shows that when gNa-soma is changed from 6 µS to 5 µS, and then to 4 µS and 3 µS, this covers the range of the AP overshoot values, as well as the maximum-rise speed values observed in recorded real-cell data during aging (see Fig. 2). Importantly, AP simulation with these gNA-soma values minimally changes the spike afterhyperpolarization amplitude and the repolarization maximum speed, and it does not effect the AP initial phase generated by Na^+^ channels in the axon (see Fig. 5B), akin to what was observed in recorded real-cell data (see Fig. 3A,B).

The simulations demonstrate that the selective reduction of high-threshold voltage-gated sodium channels located at the cell soma and proximal dendrite can mimic the selective attenuation of the AP rise speed, overshoot, and lengthening of the depolarization phase half-width akin to what was observed in real fast-spiking cells during aging.

## 4. DISCUSSION

Although various structural, metabolic, and molecular alterations have been characterized in the human brain with aging (Benavides-Piccione et al., 2013; Blazquez-Llorca et al., 2010; McQuail et al., 2021; Sibille, 2013; Ueno et al., 2018), it is still largely unknown how the electrical function of human neurons changes with age. The scarcity of functional human brain cell data is largely explained by the limited availability of living human brain tissue for research. However, in recent years, the utilization of brain tissue resected during surgery has become increasingly common, enabling the investigation of neuronal properties with demographic features (Chameh et al., 2023; Lee et al., 2020). Our study is one of the very few aging investigations (Wang et al., 2016) performed with the fast-spiking interneuron type in the human neocortex.

We found changes in the AP of these human neurons with age. These mostly showed reduced overshoot and prolonged AP depolarization phase, as well as decreased AP rise speed, even though this change was not significant in one of the three tests used. A large cell-to-cell variation in AP values measured at any age in the fast-spiking interneuron makes a large sample size, i.e., n-number, necessary for powerful statistical results. We used three types of analyses in parallel. The AP maximum rise speed significantly decreased between the youngest and oldest patient age groups, although the correlation analysis did not reach the significance level. However, because the youngest age category (“up to 30 years old”) contains cells recorded from patients in puberty, it is possible that normal developmental processes, rather than aging, affect some AP features in the data pool.

Compared to measurements and investigation of a simple AP parameter such as amplitude, these detailed AP parameters provide insights into the mechanisms underlying AP changes. These alterations indicate the attenuation of the sodium current generated at the fast-spiking neuron’s soma and proximal dendrite membrane during an AP (Bean, 2007). In fast-spiking neurons, this current is predominantly mediated by high-threshold Nav1.1-type sodium channels (Li et al., 2014; Ogiwara et al., 2007). The AP’s initial rise slope, which reflects the low-threshold sodium current generated in the axon’s initial segment (Bean, 2007; Li et al., 2014), was not changed by age. Interestingly, the AP repolarization phase was also unaffected, as shown by the unchanged spike repolarization speed and the afterhyperpolarization amplitude. This suggests that the underlying potassium current in the AP remains stable throughout a person’s lifetime. Overall, the data indicate that aging most pronouncedly affects the somatic sodium channel current of the AP.

The computational model allowed us to simulate how the “deterioration” of the high-threshold voltage-gated sodium current at the cell soma and proximal dendrite affects the AP. A single-cell computational model (Szegedi et al., 2023), which is based on realistic parameters measured in human fast-spiking neuron soma, demonstrated that selective weakening of the current at the cell soma membrane is sufficient to mimic the altered waveform and amplitude of the AP observed in real neurons by aging. Thus, computational simulation supports the idea that the high-threshold sodium channel current in cell soma is compromised in aged fast-spiking neurons.

Given that the axonal AP does not significantly change with age, it is anticipated that the neuron’s capacity to fire spikes at high frequency in the axon is not necessarily compromised. Indeed, earlier work has shown that aging is not associated with decreased maximal firing frequency of human fast-spiking neurons (Wang et al., 2016).

The somatic AP component (Bean, 2007; Li et al., 2014) generates backpropagating electrical spikes to distal dendrites in fast-spiking interneurons (Chiovini et al., 2014). In fast-spiking interneurons, AP back-propagation to dendrites is weaker than that in many other neuron types (Hu and Vervaeke, 2018). However, it is capable of activating voltage-sensitive L-type calcium channels and facilitating the activation of glutamate N-methyl-D-aspartate -receptor (NMDAR) channels in the dendrites, improving signal propagation for incoming synaptic inputs (Chiovini et al., 2014; Hu and Vervaeke, 2018; Judak et al., 2022). Weakening of the somatic sodium channel-mediated AP component in the fast-spiking interneuron soma suggests the attenuation of the AP’s back-propagation to the dendrites and reduced L-type calcium channel and glutamate NMDAR activation in dendritic branches. This sequential activation cascade of voltage-sensitive channels in dendrites occurs in fast-spiking cells during the memory consolidation process (Chiovini et al., 2014), and the deterioration of the cascade by age may be involved in reduced learning and cognitive capacity (Harada et al., 2013; Judak et al., 2022). Indeed, Nav1.1. overexpression in cortical fast-spiking interneurons is capable of restoring impaired cognitive function and weakened neuronal network activity in the neocortex in a mouse model of Alzheimer’s disease (Martinez-Losa et al., 2018). Hence, it is possible that the mechanisms underlying the AP waveform changes reported here also contribute to the senescence-related deterioration of learning and computation in the human neocortex.

Although reduced somatodendritic Na^+^ channel activity remains unclear, one possible explanation is that this alteration reduces energy consumption by fast-spiking neurons. Action potential firing consumes energy by causing partial dissipation of Na^+^ and K^+^ transmembrane ion gradients maintained by the Na^+^–K^+^ pump (Na^+^/K^+^ ATPase, sodium– potassium adenosine triphosphatase) (Attwell and Laughlin, 2001). Backpropagating action potentials with pronounced Na^+^ channel activation in dendrites increase metabolic load by increasing Na^+^/K^+^ ATPase activity (Hallermann et al., 2012). Energy metabolism is compromised in neurons in aged brains (Beishon et al., 2021), and cells may reduce their metabolic energy rate.

Our findings show that passive electrical properties (resistance, capacitance, and membrane time constant) did not significantly change in human fast-spiking cells with age. This result agrees with those of another recent study that demonstrates that these intrinsic cellular features are not changed by age in pyramidal cells of the human neocortex (Popov et al., 2023).

Given that human neurons differ in various ways from their counterparts in rodents (Chameh et al., 2023; de Kock and Feldmeyer, 2023; Eyal et al., 2016; Gidon et al., 2020; Kalmbach et al., 2018; Kim et al., 2023; Lee et al., 2023; Libe-Philippot et al., 2023; Molnar et al., 2016; Szegedi et al., 2023; Testa-Silva et al., 2022; Testa-Silva et al., 2014; Wilbers et al., 2023), it is highly relevant to study human cells directly. The life span of humans is significantly longer than that of rodents, and because neurons do not divide in the neocortex once they are generated, it is important to understand how the same cell types survive aging in humans who live 20–30 times longer than rodents do. Although human tissue heterogeneity inevitably brings cell-to-cell variation to the results (Chameh et al., 2023; Peng et al., 2019), it is noticeable that in this study, the monitoring of basic parameters through robust variables showed significant changes by age, despite the tissue source demographic variability and cortical area differences. Overall, studies conducted on neuronal properties using human resected brain tissue provide highly valuable information on how neurons function and age in the human brain.

## FIGURE LEGENDS

Table 1. Parameters for the human basket cell model.

Columns from left: Current = Transmembrane current type and conductance (g, nS) in the soma, dendrite, and axon. E (mV) = reversal potential of the current; p = exponent of the activation term.

## ACKNOWLEDGMENTS

Project no. TKP-2021-EGA-05 was implemented with the support of the Ministry of Culture and Innovation of Hungary from the National Research, Development and Innovation Fund, financed under the TKP2021-EGA funding scheme. Project no. 2022-2.1.1-NL-2022-00005 was implemented with the support of the Ministry of Culture and Innovation of Hungary from the National Research, Development and Innovation Fund, financed under the 2022-2.1.1-NL funding scheme. This project received funding from the EU’s Horizon 2020 research and innovation program under grant agreement No. 739593. This work was supported by Nemzeti Kutatási, Fejlesztési és Innovaciós Alap, OTKA K 134279, Magyar Tudományos Akadémia - the National Brain Research Program Hungary, and by University of Szeged Open Access Fund (6786). The funders played no role in the study design, data collection and analysis, decision to publish, or preparation of the manuscript.

## DECLARATION OF GENERATIVE AI AND AI-ASSISTED TECHNOLOGIES IN THE WRITING PROCESS

During the preparation of this work the authors used Enago editing service in order to improve the English language of this manuscript. After using this service, the authors reviewed and edited the content as needed and take full responsibility for the content of the publication.

## AUTHOR CONTRIBUTIONS

VS: Visualization, Data Curation, Investigation, Formal analysis AT: Data Curation, Formal analysis SF: Investigation AD: Investigation EB: Investigation PB: Resources GT: Resources AS: Visualization, Software, Methodology KL: Funding acquisition, Supervision, Writing - Review & Editing, Investigation

## REFERENCES

Aranda-Anzaldo, A., Dent, M.A., (2017) Why Cortical Neurons Cannot Divide, and Why Do They Usually Die in the Attempt? J Neurosci Res 95, 921–929.

Bean, B.P., (2007) The action potential in mammalian central neurons. Nat Rev Neurosci 8, 451–465.

Attwell, D., Laughlin, S.B., (2001) An energy budget for signaling in the grey matter of the brain. J Cereb Blood Flow Metab 21: 1133–1145.

Beishon, L., Clough, R.H., Kadicheeni, M., Chithiramohan, T., Panerai, R.B., Haunton, V.J., Minhas, J.S., Robinson, T.G., (2021) Vascular and haemodynamic issues of brain ageing. Pflugers Arch 473, 735–751.

Benavides-Piccione, R., Fernaud-Espinosa, I., Robles, V., Yuste, R., DeFelipe, J., (2013) Age-based comparison of human dendritic spine structure using complete three-dimensional reconstructions. Cereb Cortex 23, 1798–1810.

Blazquez-Llorca, L., Garcia-Marin, V., DeFelipe, J., (2010) GABAergic complex basket formations in the human neocortex. J Comp Neurol 518, 4917–4937.

Cernotova, D., Hruzova, K., Levcik, D., Svoboda, J., Stuchlik, A., (2023) Linking Social Cognition, Parvalbumin Interneurons, and Oxytocin in Alzheimer’s Disease: An Update. J Alzheimers Dis 96, 861–875.

Chameh, H.M., Falby, M., Movahed, M., Arbabi, K., Rich, S., Zhang, L., Lefebvre, J., Tripathy, S.J., De Pitta, M., Valiante, T.A., (2023) Distinctive biophysical features of human cell-types: insights from studies of neurosurgically resected brain tissue. Front Synaptic Neurosci 15, 1250834.

Chiovini, B., Turi, G.F., Katona, G., Kaszas, A., Palfi, D., Maak, P., Szalay, G., Szabo, M.F., Szabo, G., Szadai, Z., Kali, S., Rozsa, B., (2014) Dendritic spikes induce ripples in parvalbumin interneurons during hippocampal sharp waves. Neuron 82, 908–924.

Cunnane, S.C., Trushina, E., Morland, C., Prigione, A., Casadesus, G., Andrews, Z.B., Beal, M.F., Bergersen, L.H., Brinton, R.D., de la Monte, S., Eckert, A., Harvey, J., Jeggo, R., Jhamandas, J.H., Kann, O., la Cour, C.M., Martin, W.F., Mithieux, G., Moreira, P.I., Murphy, M.P., Nave, K.A., Nuriel, T., Oliet, S.H.R., Saudou, F., Mattson, M.P., Swerdlow, R.H., Millan, M.J., (2020) Brain energy rescue: an emerging therapeutic concept for neurodegenerative disorders of ageing. Nat Rev Drug Discov 19, 609–633.

de Kock, C.P.J., Feldmeyer, D., (2023) Shared and divergent principles of synaptic transmission between cortical excitatory neurons in rodent and human brain. Front Synaptic Neurosci 15, 1274383.

Dong, Y., Zhao, K., Qin, X., Du, G., Gao, L., (2023) The mechanisms of perineuronal net abnormalities in contributing aging and neurological diseases. Ageing Res Rev 92, 102092.

Druga, R., Salaj, M., Al-Redouan, A., (2023) Parvalbumin - Positive Neurons in the Neocortex: A Review. Physiol Res 72, S173–S191.

Eyal, G., Verhoog, M.B., Testa-Silva, G., Deitcher, Y., Lodder, J.C., Benavides-Piccione, R., Morales, J., DeFelipe, J., de Kock, C.P., Mansvelder, H.D., Segev, I., (2016) Unique membrane properties and enhanced signal processing in human neocortical neurons. Elife 5.

Gidon, A., Zolnik, T.A., Fidzinski, P., Bolduan, F., Papoutsi, A., Poirazi, P., Holtkamp, M., Vida, I., Larkum, M.E., (2020) Dendritic action potentials and computation in human layer 2/3 cortical neurons. Science 367, 83–87.

Guet-McCreight, A., Chameh, H.M., Mahallati, S., Wishart, M., Tripathy, S.J., Valiante, T.A., Hay, E., (2023) Age-dependent increased sag amplitude in human pyramidal neurons dampens baseline cortical activity. Cereb Cortex 33, 4360–4373.

Hallermann, S., de Kock, C. P. J., Stuart, G. J., Kole, M. H., (2012) State and location dependence of action potential metabolic cost in cortical pyramidal neurons. Nature Neuroscience, 15(7), 1007–1014. 10.1038/nn.3132

Harada, C.N., Natelson Love, M.C., Triebel, K.L., (2013) Normal cognitive aging. Clin Geriatr Med 29, 737–752.

Herrup, K., (2013) Post-mitotic role of the cell cycle machinery. Curr Opin Cell Biol 25, 711–716.

Higley, M.J., Cardin, J.A., (2022) Spatiotemporal dynamics in large-scale cortical networks. Curr Opin Neurobiol 77, 102627.

Hijazi, S., Smit, A.B., van Kesteren, R.E., (2023) Fast-spiking parvalbumin-positive interneurons in brain physiology and Alzheimer’s disease. Mol Psychiatry. 10.1038/s41380-023-02168-y

Hu, H., Gan, J., Jonas, P., (2014) Interneurons. Fast-spiking, parvalbumin(+) GABAergic interneurons: from cellular design to microcircuit function. Science 345, 1255263.

Hu, H., Vervaeke, K., (2018) Synaptic integration in cortical inhibitory neuron dendrites. Neuroscience 368, 115–131.

Ibrahim, B.A., Llano, D.A., (2019) Aging and Central Auditory Disinhibition: Is It a Reflection of Homeostatic Downregulation or Metabolic Vulnerability? Brain Sci 9.

Isaev, N.K., Stelmashook, E.V., Genrikhs, E.E., (2019) Neurogenesis and brain aging. Rev Neurosci 30, 573–580.

Judak, L., Chiovini, B., Juhasz, G., Palfi, D., Mezriczky, Z., Szadai, Z., Katona, G., Szmola, B., Ocsai, K., Martinecz, B., Mihaly, A., Denes, A., Kerekes, B., Szepesi, A., Szalay, G., Ulbert, I., Mucsi, Z., Roska, B., Rozsa, B., (2022) Sharp-wave ripple doublets induce complex dendritic spikes in parvalbumin interneurons in vivo. Nat Commun 13, 6715.

Kalmbach, B.E., Buchin, A., Long, B., Close, J., Nandi, A., Miller, J.A., Bakken, T.E., Hodge, R.D., Chong, P., de Frates, R., Dai, K., Maltzer, Z., Nicovich, P.R., Keene, C.D., Silbergeld, D.L., Gwinn, R.P., Cobbs, C., Ko, A.L., Ojemann, J.G., Koch, C., Anastassiou, C.A., Lein, E.S., Ting, J.T., (2018) h-Channels Contribute to Divergent Intrinsic Membrane Properties of Supragranular Pyramidal Neurons in Human versus Mouse Cerebral Cortex. Neuron 100, 1194–1208 e1195.

Kann, O., (2016) The interneuron energy hypothesis: Implications for brain disease. Neurobiol Dis 90, 75–85.

Kim, M.H., Radaelli, C., Thomsen, E.R., Monet, D., Chartrand, T., Jorstad, N.L., Mahoney, J.T., Taormina, M.J., Long, B., Baker, K., Bakken, T.E., Campagnola, L., Casper, T., Clark, M., Dee, N., D’Orazi, F., Gamlin, C., Kalmbach, B.E., Kebede, S., Lee, B.R., Ng, L., Trinh, J., Cobbs, C., Gwinn, R.P., Keene, C.D., Ko, A.L., Ojemann, J.G., Silbergeld, D.L., Sorensen, S.A., Berg, J., Smith, K.A., Nicovich, P.R., Jarsky, T., Zeng, H., Ting, J.T., Levi, B.P., Lein, E., (2023) Target cell-specific synaptic dynamics of excitatory to inhibitory neuron connections in supragranular layers of human neocortex. Elife 12.

Lee, B.R., Dalley, R., Miller, J.A., Chartrand, T., Close, J., Mann, R., Mukora, A., Ng, L., Alfiler, L., Baker, K., Bertagnolli, D., Brouner, K., Casper, T., Csajbok, E., Donadio, N., Driessens, S.L.W., Egdorf, T., Enstrom, R., Galakhova, A.A., Gary, A., Gelfand, E., Goldy, J., Hadley, K., Heistek, T.S., Hill, D., Hou, W.H., Johansen, N., Jorstad, N., Kim, L., Kocsis, A.K., Kruse, L., Kunst, M., Leon, G., Long, B., Mallory, M., Maxwell, M., McGraw, M., McMillen, D., Melief, E.J., Molnar, G., Mortrud, M.T., Newman, D., Nyhus, J., Opitz-Araya, X., Ozsvar, A., Pham, T., Pom, A., Potekhina, L., Rajanbabu, R., Ruiz, A., Sunkin, S.M., Szots, I., Taskin, N., Thyagarajan, B., Tieu, M., Trinh, J., Vargas, S., Vumbaco, D., Waleboer, F., Walling-Bell, S., Weed, N., Williams, G., Wilson, J., Yao, S., Zhou, T., Barzo, P., Bakken, T., Cobbs, C., Dee, N., Ellenbogen, R.G., Esposito, L., Ferreira, M., Gouwens, N.W., Grannan, B., Gwinn, R.P., Hauptman, J.S., Hodge, R., Jarsky, T., Keene, C.D., Ko, A.L., Korshoej, A.R., Levi, B.P., Meier, K., Ojemann, J.G., Patel, A., Ruzevick, J., Silbergeld, D.L., Smith, K., Sorensen, J.C., Waters, J., Zeng, H., Berg, J., Capogna, M., Goriounova, N.A., Kalmbach, B., de Kock, C.P.J., Mansvelder, H.D., Sorensen, S.A., Tamas, G., Lein, E.S., Ting, J.T., (2023) Signature morphoelectric properties of diverse GABAergic interneurons in the human neocortex. Science 382, eadf6484.

Lee, K., Park, T.I., Heppner, P., Schweder, P., Mee, E.W., Dragunow, M., Montgomery, J.M., (2020) Human in vitro systems for examining synaptic function and plasticity in the brain. J Neurophysiol 123, 945–965.

Lehmann, K., Steinecke, A., Bolz, J., (2012) GABA through the ages: regulation of cortical function and plasticity by inhibitory interneurons. Neural Plast 2012, 892784.

Li, T., Tian, C., Scalmani, P., Frassoni, C., Mantegazza, M., Wang, Y., Yang, M., Wu, S., Shu, Y., (2014) Action potential initiation in neocortical inhibitory interneurons. PLoS Biol 12, e1001944.

Libe-Philippot, B., Lejeune, A., Wierda, K., Louros, N., Erkol, E., Vlaeminck, I., Beckers, S., Gaspariunaite, V., Bilheu, A., Konstantoulea, K., Nyitrai, H., De Vleeschouwer, M., Vennekens, K.M., Vidal, N., Bird, T.W., Soto, D.C., Jaspers, T., Dewilde, M., Dennis, M.Y., Rousseau, F., Comoletti, D., Schymkowitz, J., Theys, T., de Wit, J., Vanderhaeghen, P., (2023) LRRC37B is a human modifier of voltage-gated sodium channels and axon excitability in cortical neurons. Cell 186, 5766–5783 e5725.

Liu, Y., Konopka, G., (2020) An integrative understanding of comparative cognition: lessons from human brain evolution. Integr Comp Biol 60, 991–1006.

Luhmann, H.J., (2023) Dynamics of neocortical networks: connectivity beyond the canonical microcircuit. Pflugers Arch 475, 1027–1033.

Martinez-Losa, M., Tracy, T.E., Ma, K., Verret, L., Clemente-Perez, A., Khan, A.S., Cobos, I., Ho, K., Gan, L., Mucke, L., Alvarez-Dolado, M., Palop, J.J., (2018) Nav1.1-Overexpressing Interneuron Transplants Restore Brain Rhythms and Cognition in a Mouse Model of Alzheimer’s Disease. Neuron 98, 75–89 e75.

Mattson, M.P., (2020) Involvement of GABAergic interneuron dysfunction and neuronal network hyperexcitability in Alzheimer’s disease: Amelioration by metabolic switching. Int Rev Neurobiol 154, 191–205.

McQuail, J.A., Beas, B.S., Kelly, K.B., Hernandez, C.M., 3rd, Bizon, J.L., Frazier, C.J., (2021) Attenuated NMDAR signaling on fast-spiking interneurons in prefrontal cortex contributes to age-related decline of cognitive flexibility. Neuropharmacology 197, 108720.

Molnar, G., Rozsa, M., Baka, J., Holderith, N., Barzo, P., Nusser, Z., Tamas, G., (2016) Human pyramidal to interneuron synapses are mediated by multi-vesicular release and multiple docked vesicles. Elife 5.

Ogiwara, I., Miyamoto, H., Morita, N., Atapour, N., Mazaki, E., Inoue, I., Takeuchi, T., Itohara, S., Yanagawa, Y., Obata, K., Furuichi, T., Hensch, T.K., Yamakawa, K., (2007) Nav1.1 localizes to axons of parvalbumin-positive inhibitory interneurons: a circuit basis for epileptic seizures in mice carrying an Scn1a gene mutation. J Neurosci 27, 5903–5914.

Pegasiou, C.M., Zolnourian, A., Gomez-Nicola, D., Deinhardt, K., Nicoll, J.A.R., Ahmed, A.I., Vajramani, G., Grundy, P., Verhoog, M.B., Mansvelder, H.D., Perry, V.H., Bulters, D., Vargas-Caballero, M., (2020) Age-Dependent Changes in Synaptic NMDA Receptor Composition in Adult Human Cortical Neurons. Cereb Cortex 30, 4246–4256.

Peng, Y., Mittermaier, F.X., Planert, H., Schneider, U.C., Alle, H., Geiger, J.R.P., (2019) High-throughput microcircuit analysis of individual human brains through next-generation multineuron patch-clamp. Elife 8.

Peters, R., (2006) Ageing and the brain. Postgrad Med J 82, 84–88.

Petilla Interneuron Nomenclature, G., Ascoli, G.A., Alonso-Nanclares, L., Anderson, S.A., Barrionuevo, G., Benavides-Piccione, R., Burkhalter, A., Buzsaki, G., Cauli, B., Defelipe, J., Fairen, A., Feldmeyer, D., Fishell, G., Fregnac, Y., Freund, T.F., Gardner, D., Gardner, E.P., Goldberg, J.H., Helmstaedter, M., Hestrin, S., Karube, F., Kisvarday, Z.F., Lambolez, B., Lewis, D.A., Marin, O., Markram, H., Munoz, A., Packer, A., Petersen, C.C., Rockland, K.S., Rossier, J., Rudy, B., Somogyi, P., Staiger, J.F., Tamas, G., Thomson, A.M., Toledo-Rodriguez, M., Wang, Y., West, D.C., Yuste, R., (2008) Petilla terminology: nomenclature of features of GABAergic interneurons of the cerebral cortex. Nat Rev Neurosci 9, 557–568.

Popov, A., Brazhe, N., Morozova, K., Yashin, K., Bychkov, M., Nosova, O., Sutyagina, O., Brazhe, A., Parshina, E., Li, L., Medyanik, I., Korzhevskii, D.E., Shenkarev, Z., Lyukmanova, E., Verkhratsky, A., Semyanov, A., (2023) Mitochondrial malfunction and atrophy of astrocytes in the aged human cerebral cortex. Nat Commun 14, 8380.

Radulescu, C.I., Cerar, V., Haslehurst, P., Kopanitsa, M., Barnes, S.J., (2021) The aging mouse brain: cognition, connectivity and calcium. Cell Calcium 94, 102358.

Rozycka, A., Liguz-Lecznar, M., (2017) The space where aging acts: focus on the GABAergic synapse. Aging Cell 16, 634–643.

Sibille, E., (2013) Molecular aging of the brain, neuroplasticity, and vulnerability to depression and other brain-related disorders. Dialogues Clin Neurosci 15, 53–65.

Straehle, J., Ravi, V.M., Heiland, D.H., Galanis, C., Lenz, M., Zhang, J., Neidert, N.N., El Rahal, A., Vasilikos, I., Kellmeyer, P., Scheiwe, C., Klingler, J.H., Fung, C., Vlachos, A., Beck, J., Schnell, O., (2023) Technical report: surgical preparation of human brain tissue for clinical and basic research. Acta Neurochir (Wien) 165, 1461–1471.

Szegedi, V., Bakos, E., Furdan, S., Kovacs, B.H., Varga, D., Erdelyi, M., Barzo, P., Szucs, A., Tamas, G., Lamsa, K., (2023) HCN channels at the cell soma ensure the rapid electrical reactivity of fast-spiking interneurons in human neocortex. PLoS Biol 21, e3002001.

Temple, S., (2023) Advancing cell therapy for neurodegenerative diseases. Cell Stem Cell 30, 512–529.

Testa-Silva, G., Rosier, M., Honnuraiah, S., Guzulaitis, R., Megias, A.M., French, C., King, J., Drummond, K., Palmer, L.M., Stuart, G.J., (2022) High synaptic threshold for dendritic NMDA spike generation in human layer 2/3 pyramidal neurons. Cell Rep 41, 111787.

Testa-Silva, G., Verhoog, M.B., Linaro, D., de Kock, C.P., Baayen, J.C., Meredith, R.M., De Zeeuw, C.I., Giugliano, M., Mansvelder, H.D., (2014) High bandwidth synaptic communication and frequency tracking in human neocortex. PLoS Biol 12, e1002007.

Ueno, H., Takao, K., Suemitsu, S., Murakami, S., Kitamura, N., Wani, K., Okamoto, M., Aoki, S., Ishihara, T., (2018) Age-dependent and region-specific alteration of parvalbumin neurons and perineuronal nets in the mouse cerebral cortex. Neurochem Int 112, 59–70.

Wang, B., Ke, W., Guang, J., Chen, G., Yin, L., Deng, S., He, Q., Liu, Y., He, T., Zheng, R., Jiang, Y., Zhang, X., Li, T., Luan, G., Lu, H.D., Zhang, M., Zhang, X., Shu, Y., (2016) Firing Frequency Maxima of Fast-Spiking Neurons in Human, Monkey, and Mouse Neocortex. Front Cell Neurosci 10, 239.

Wilbers, R., Metodieva, V.D., Duverdin, S., Heyer, D.B., Galakhova, A.A., Mertens, E.J., Versluis, T.D., Baayen, J.C., Idema, S., Noske, D.P., Verburg, N., Willemse, R.B., de Witt Hamer, P.C., Kole, M.H.P., de Kock, C.P.J., Mansvelder, H.D., Goriounova, N.A., (2023) Human voltagegated Na(+) and K(+) channel properties underlie sustained fast AP signaling. Sci Adv 9, eade3300.

