## Supplemental table 1 for "Aging-associated weakening of the action potential in fast-spiking interneurons in the human neocortex"

| Current | *g_soma_* | *g_den_* | *g_ax_* | *E* | *p* | *V_m,1/2_* | *V_m,sl_* | *V_h,1/2_* | *V_h,sl_* | *τ_m,max_* | *τ_m,min_* | *V_tm,1/2_* | *V_tm,sl_* | *τ_h,max_* | *τ_h,min_* | *V_th,1/2_* | *V_th,sl_* |
| --- | --- | --- | --- | --- | --- | --- | --- | --- | --- | --- | --- | --- | --- | --- | --- | --- | --- |
|  | nS | nS | nS | mV |  | mV | mV | mV | mV | ms | ms | mV | mV | ms | ms | mV | mV |
| Na1.1 | 6000 |  | 3500 | 50 | 3 | -26 | 10 | -62 | -17 | 0.7 | 0.1 | -68 | 30 | 4 | 0.2 | -74 | 30 |
| Na1.6 |  |  | 1130 | 50 | 3 | -44 | 10 | -82 | -18 | 0.7 | 0.1 | -68 | 30 | 4 | 0.2 | -74 | 30 |
| HCN | 0.3 | 0.4 |  | -30 | 1 | -73 | -16 |  |  | 200 | 100 | -45 | 50 |  |  |  |  |
| K_d_ | 350 |  | 350 | -75 | 4 | -26 | 15 |  |  | 2 | 0.5 | -70 | 30 |  |  |  |  |
| K_ir_ | 1.0 | 0.6 |  | -75 | 1 | -88 | -12 |  |  | 10 | 1.0 | -45 | 35 |  |  |  |  |
| D | 420 |  | 420 | -75 | 1 | -41 | 17 | -68 | -13 | 4 | 1 | -80 | 80 | 340 | 84 | -90 | 70 |
